# Large-scale causal discovery using interventional data sheds light on the regulatory network architecture of blood traits

**DOI:** 10.1101/2023.10.13.562293

**Authors:** Brielin C. Brown, John A. Morris, Tuuli Lappalainen, David A. Knowles

## Abstract

Inference of directed biological networks is an important but notoriously challenging problem. We introduce *inverse* sparse *regression (inspre)*, an approach to learning causal networks that leverages large-scale intervention-response data. Applied to 788 genes from the genome-wide perturb-seq dataset, *inspre* helps elucidate the network architecture of blood traits.

## Main

Recent developments in the understanding of complex-trait genetics have lead to a call for increased study of directed biological networks because they are crucial for dissecting the genetic architecture of complex traits and finding pathways that can be targeted for treatment^1–3^. However, interrogating causal structure (i.e. *causal discovery*) is notoriously difficult owing to factors such as unmeasured confounding, reverse causation and the presence of cycles^4^. Even assuming that all relevant variables are measured and that the underlying graph is acyclic, the exact network is still not identifiable using observational data alone^5^, and identifying the set of observationally-equivalent graphs is computationally intractable^6^.

Interventional data improves the identifiability of causal models^7^ and can eliminate biases due to unobserved confounding^8^. Therefore, advances in the scope and scale of gene-targeting CRISPR-intervention experiments^9^ have created an ideal setting for the development of methods for large-scale causal graph inference. Recent advancements include dotears^10^, which extends the convex optimization-based approach notears^11^ to interventional data, and igsp^12^, which learns equivalence classes of graphs using a permutation based approach. However, existing methods can assume a strong intervention model^10^, return an unweighted graph^12^ or be intractable for large graphs^10,11^. They also generally assume the graph is acyclic and unconfounded^10–12^.

Here, we introduce an approach to causal discovery from interventional data using a two-stage procedure (Figure 1a). After estimating the marginal average causal effect (ACE) of every feature on every other, the matrix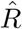, the causal graph *G* can be obtained via the relationship *G* = *I − R*^*−*1^*D*[1*/R*^*−*1^], where */* indicates element-wise division and the operator *D*[*A*] sets off-diagonal entries of the matrix to 0^13^. Importantly, while *G* can be determined exactly given the true matrix *R*, we only have access to a noisy estimate 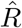, which may not be well-conditioned or even invertible. Our primary contribution is a procedure for estimating a *sparse approximate inverse* of the ACE matrix via solving the constrained optimization problem

**Figure 1:**
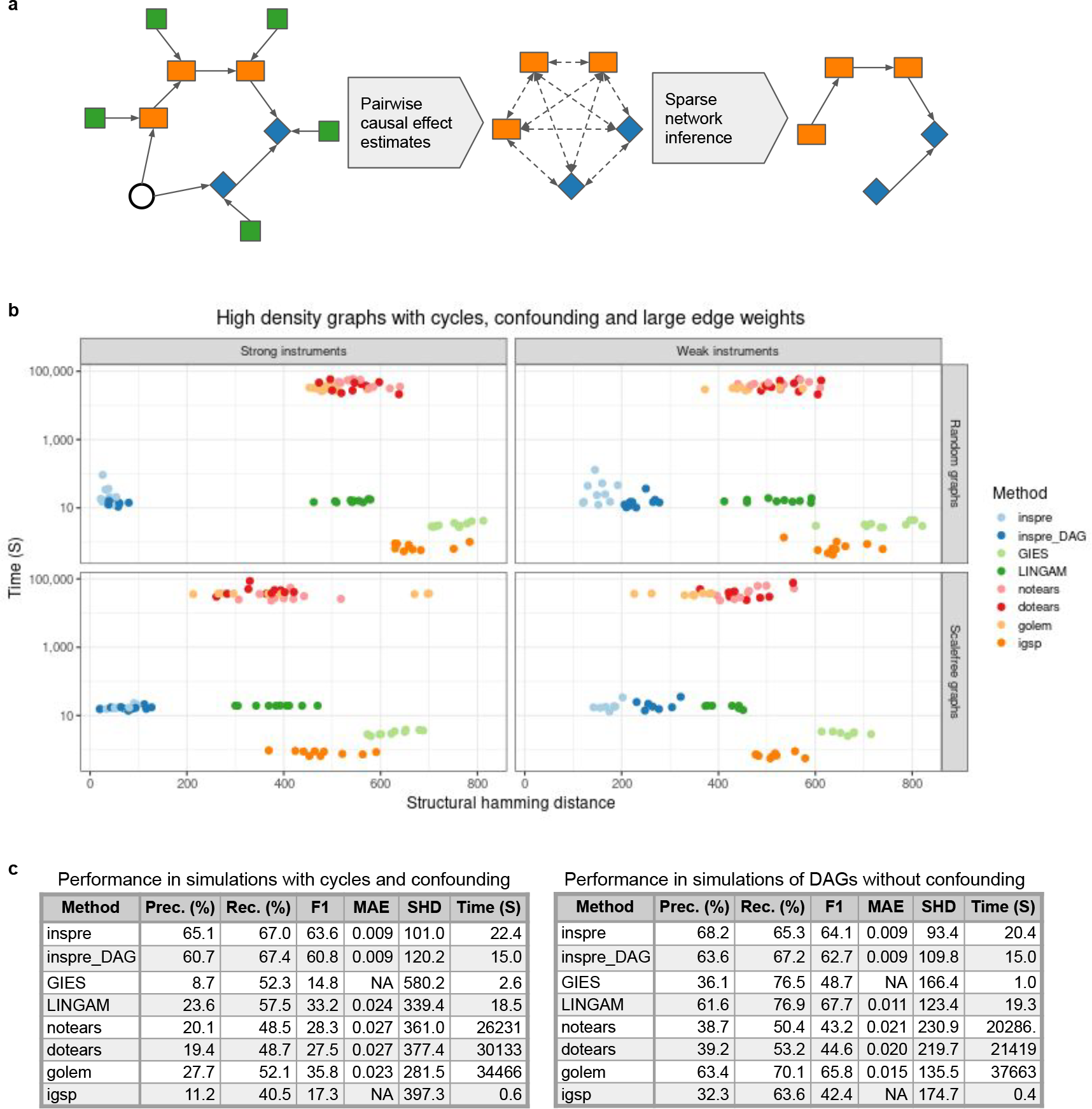
a) Overview of the *inspre* approach. Interventions (green squares) are used to estimate marginal average causal effects between all pairs of features (orange rectangles and blue diamonds). These pairwise estimates are used to infer a sparse network on the features. b) Structural hamming distance versus time for *inspre* compared to commonly-used methods in 5the setting of high density graphs with confounding, cycles, and large edge weights (lower is better). inspre DAG represents *inspre* with an approximate acyclicity constriant, see Methods for details. c) Averaged over density, intervention strength, graph type, edge effect size *inspre* is the most performant approach by several metrics even in acyclic graphs without confounding.

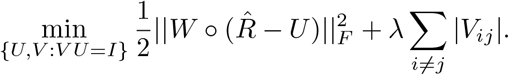

This approximate inverse is then used to estimate *G* via *Ĝ* = *I − V D*[1*/V*]. We call this approach *inverse sparse regression (inspre)*. Here, *U* approximates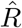 while its left-inverse *V* has sparsity controlled via the *L*_1_ optimization parameter *λ*. The weight matrix *W* allows us to place less emphasis on entries of 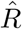 with high standard error. For complete details see Methods.

Working with the bi-directional ACE matrix *R* rather than the full data matrix provides several advantages. First, interventional data can be used to estimate effects that are robust to unobserved confounding. Second, leveraging bi-directed ACE estimates that include both the effect of feature *i* on *j* and *j* on *i* allows us to accommodate graphs with cycles. Finally, the features by features ACE matrix is typically much smaller than than the original samples by features data matrix, providing a dramatic speedup that enables inference in settings with hundreds or even thousands of features.

We evaluated the performance of *inspre* under a host of different simulation settings while comparing against other commonly-used methods for causal discovery from observational (LinGAM^14^, notears^11^, golem^15^) and interventional (GIES^16^, igsp^12^, dotears^10^) data. We tested 50-node cyclic and acyclic graphs with 100 interventional samples per node and 5000 total control samples. We simulated graphs with and without confounding while varying the graph type (Erd ő s-Réyni random^17^ vs scale-free^18^), density (high vs low), edge weights (large vs small), and intervention strength (strong vs weak). We conducted each of these 64 experiments 10 times and compared methods by structural Hamming distance (SHD), precision, recall, F1-score, mean absolute error and runtime (Figure 1b-c, Table S1).

*inspre* out-performs other tested methods in cyclic graphs with confounding by a large margin, even when interventions are weak (Figure 1b). Perhaps more interestingly, *inspre* still obtains the highest precision, lowest SHD and lowest MAE in acyclic graphs without confounding when averaged over graph type, density, edge weight and intervention strength (Figure 1c). On top of this, *inspre* takes just seconds to run, while comparable optimization-based approaches can take up to 10 hours. Of course, the performance of *inspre* is dependent on edge weight and intervention strength when network effects are small and interventions are weak, *inspre* performs comparatively poorly. However even in this setting, the weighting scheme biases our approach to high precision, low recall, and the resulting SHD is still comparable to other methods even in the absence of confounding (Table S1).

We applied *inspre* to the K562 genome-wide Perturb-seq (GWPS)^9^ experiment targeting essential genes. We selected 788 genes on the basis of guide effectiveness and number of cells receiving a guide targeting that gene. After estimating the ACE of each gene on every other, we found 131,155 significant effects at FDR 5%. We then used *inspre* to construct a graph on these 788 nodes, which contained 10,423 edges (1.68% non-zero, Figure 2a). We used this graph to calculate shortest paths and the total effect induced by this path for all pairs of vertices. 30.9% of gene pairs are connected by at least one path, with a median path length of 2.9 (standard deviation *sd* = 0.86) for all pairs and 2.68 (*sd* = 0.83) for FDR significant pairs (Figure 2b). We then calculated the percentage of the total effect explained by the shortest path for all FDR 5% significant gene pairs. If this number is low, there are many paths between a given pair of nodes with the shortest making up only a fraction of the total effect; if this number is close to 1, the shortest path explains most of the total effect; if this number is above 1, other paths in the network work to cancel out some of the effect of the shortest path. We find that the average effect explained by the shortest path is low (median = 10.77%), and that there are many pairs where the effect explained exceeds 100% (4, 183 pairs, Figure 2c). This indicates there tends to be many important network paths when considering the effect of one gene on another.

**Figure 2:**
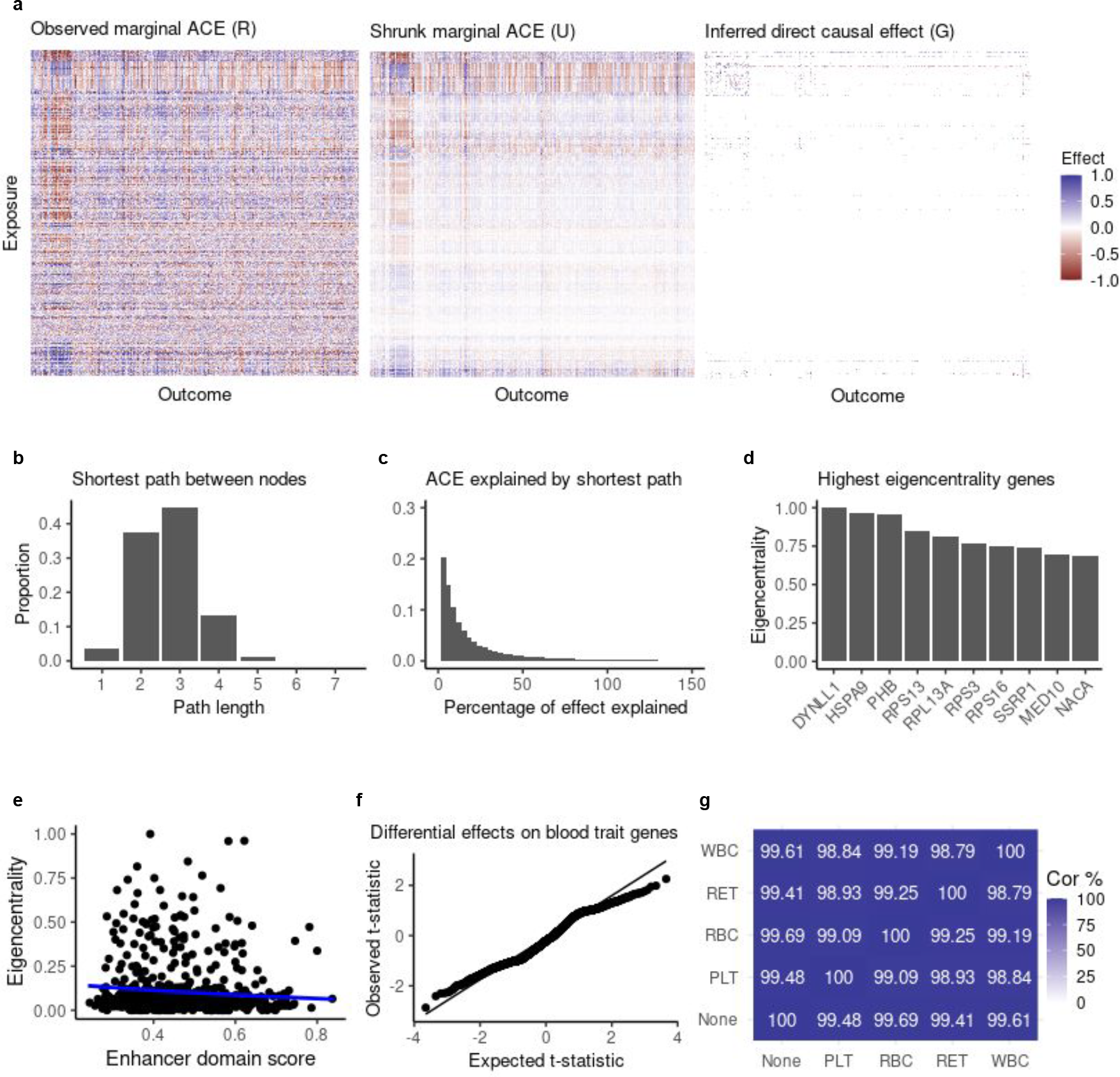
a) For 788 genes from the genome-wide perturb-seq experiment, we calculate pairwise marginal average causal effects (left). Using *inspre*, we infer a shrunk-approximation to this matrix that reduces noise (center) and is supported by an estimate of the underlying causal graph (right). b) The shortest path between pairs of nodes which have an FDR 5% significant total effect. c) The percentage of the total effect explained by the shortest path indicates that many paths are responsible for the total effect of one gene on another. d) The highest eigencentrality genes. e) Eigenentrality 6within our network is related to gene properties from orthogonal data sources, enhancer domain score is shown here (see also Figure S1, Table S4). f) QQ-plot for t-tests of a difference in mean effect of each gene on genes that have been associated with different types of blood traits in GWAS. We observe no deviation from expected. g) Correlation of the average absolute effect of each gene on those that have been associated with blood traits. Effects across categories are nearly identical.

We calculated the eigencentrality^19^ for each of the 788 nodes in our network. The most central genes include DYNLL1 (dynein light chain 1), HSPA9 (heat shock 70 kDa protein 9), PHB (prohibitin), MED10 (mediator complex subunit 10) and NACA (Nascent-polypeptide-associated complex alpha polypeptide, Figure 2c). These are highly conserved genes that play important roles in key cellular processes, particularly transcriptional regulation^20–23^. Top central genes also include several ribosomal proteins: RPS13, RRPL13A, RPS3, and RPS16 (Figure 2c). We next asked whether centrality was related to other measures of gene importance from sources such as gnomAD^24^ and ExAC^25^. We fit a beta regression model of eigencentrality on 16 such annotations while controlling the family-wise error rate using Holm’s method^26^ (Table S2). We found a strong negative association between eigencentrality and enhancer domain score^27^ (*p*_*adj*_ = 6.4 *×* 10^*−*6^, Figure 2e), indicating that genes that are more dosage sensitive also tend to be less central within the network. We also found a positive associations between eigencentrality and number of protein-protein interactions^28^(*p*_*adj*_ = 1.7 *×* 10^*−*3^, Figure S1a), gnomAD loss-of-function constrained genes (*p*_*adj*_ = 0.023, Figure S1b), and ExAC deletion intolerance (*p*_*adj*_ = 0.049, Figure S1c). We found weaker associations with gnomAD missense-constrained genes, haploinsufficiency index^29^ and CCDG deletion sensitivity^30^ (*p*_*adj*_ = 0.07 for each association, Figure S1d-f).

Finally, we sought to determine whether genes associated with classes of blood traits clustered together in pathways or were broadly distributed throughout the network. As in our previous work^31^, we used finemapping of UK Biobank^32^ phenotypes to obtain a list of genes associated with platelet (PLT), reticulocyte (RET), red blood cell (RBC), and white blood cell (WBC) traits (Figure S2, Table S2). We calculated the mean distance between genes within each trait class and tested whether this was significantly different from the distance between genes across trait categories. We found small but statistically significant differences in mean path length between versus across most blood traits (Figure 2e, Table S4). For example, the mean path length within WBC associated genes was 2.81 (standard error *se* = 0.007) while the mean path length between WBC and PLT genes was 2.93 (*se* = 0.006, *p*_*adj*_ = 1*×* 10^*−*11^), and the mean path length between WBC and RBC genes was 2.89 (*se* = 0.005, *p*_*adj*_ = 2.4*×* 10^*−*5^).

Given this, we wondered whether the network could be used to identify upstream regulators that differentiate between blood traits. For each of our 788 genes, we calculated the mean absolute effect on genes in each trait class and tested if this effect was larger than on the genes in the other trait classes. Interestingly, we found no statistically significant differences in mean absolute effect of any gene on GWAS genes for any trait class compared to the others (Figure 2f, Table S5). Instead, upstream effects on blood trait GWAS genes were nearly identical regardless of trait type (Figure 2g, Table S6).

At face value, our results support the notion that blood-trait-relevant gene networks are sufficiently connected that perturbations to most genes have broad downstream effects, rather than having effects that concentrate in particular trait-relevant pathways^1,3^. However, there are several caveats that limit the interpretability of these results. First, while K562 cells are commonly-used in CRISPR-inhibition experiments to model blood traits^31,33^, it is unclear what the relationships are between the regulatory networks in this cancer cell line and diverse blood cell populations. Along these lines, it is unclear whether the network connections discovered via said experiments are the same as those that the molecular effects of GWAS variants propagate along. Instead, large-scale gene promoter inhibition may result in changes to cellular state that brings a concomitant change in network structure. In addition, we are limited to analyzing 788 genes that were well-captured in this screen. While other genes were targeted, many of them were not captured at sufficient levels in control samples to obtain robust estimates of the intervention effect and thus were removed from our analysis. Including more genes may improve our ability to identify pathway-specific rather than broadly acting effects. Finally, when integrating with blood trait GWAS genes we analyzed sets of genes related to classes of blood traits, rather than individual ones. This may smooth-over trait-specific differences yielding a bias towards discovering broad rather than pathway specific effects.

*inspre* leverages the ACE matrix to rapidly produce causal graph estimates even with cyclic graph structures and the presence of unmeasured confounding, enabling causal discovery at unprecedented scales. The utility of *inspre* is two-fold: it provides an estimate of the causal graph while also providing a shrunken estimate of the ACE matrix that is supported by a graph structure, thereby substantially denoising the data. However, it does have some limitations. First, when intervention effects are weak, *inspre* is biased towards producing sparse, high-precision, low-recall estimates. Second, *inspre* requires an intervention on every feature and cannot make use of observed data for features without interventions. Finally, *inspre* produces an approximate inverse that may result in a graph that does not satisfy the spectral radius requirement (see Methods).

This work represents a first step towards integrating high-dimensional causal networks from CRISPR-inhibition experiments with complex human phenotypes. In the future, we anticipate improvements in this methodology combined with simultaneous increases in the scope and scale of diverse molecular intervention experiments will enable new insights into the causal structures underlying complex human disease.

## Methods

### Data model

We utilize the common linear autoregressive graph model^11^ while adding linear intervention effects. *Y* be an *N× D* matrix of features (genes, phenotypes) and *G* a *D× D* graph acting on the features in the form of a weighted adjacency matrix. Let *X* be an *N× Q* binary indicator matrix of intervention targets with *Q× D* effect matrix *β*. The model is

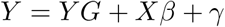

where *γ* is a mean 0 random effect representing unmeasured factors and noise. Our goal is to estimate *G* given *Y* and *X*. Note that in general, the covariance matrix of *γ* need not be isotropic. This will occur for example in the case of unmeasured confounding. Note also that in order for this model to be well defined, we require that the modulus of the largest eigenvalue of *G* (the “spectral radius” *r*(*G*)) to be less than 1. In this case, *I − G* is invertible and the model can be rewritten as

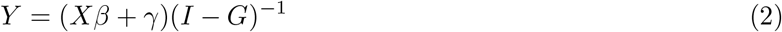

Our approach to estimating *G* is to use the interventional data to first estimate the *D× D* matrix of average causal effects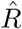 (as in 𝔼 [*Y*_*j*_ *do*(*Y*_*i*_)] = *R*_*ij*_*Y*_*i*_). We then use this matrix to estimate *G*. In practice, any approach that is consistent given the assumptions of the data setting can be used to estimate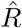. For simplicity in this derivation, we assume that there is one intervention on each feature and that these interventions are uncorrelated with each other. In this case, we can use two-stage least squares. The entries on the diagonal of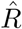 are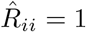. The off-diagonal elements are given by

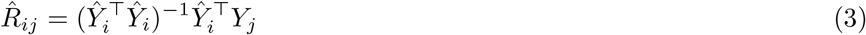

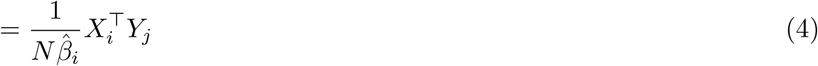

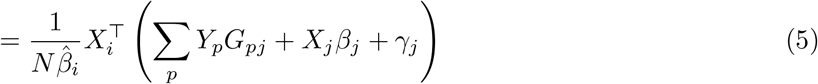

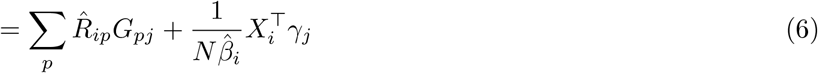

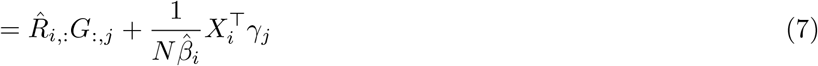

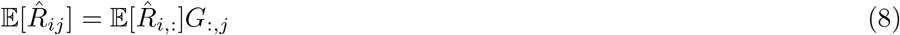

Since 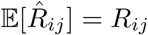, we have that *R* satisfies the recurrence *R* = *RG* off the diagonal, from which it follows that^13^,

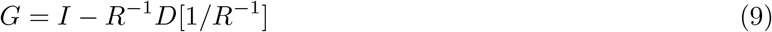

where */* indicates elementwise division and the diagonal operator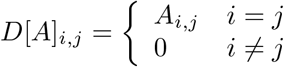 sets off-diagonal elements of a matrix to 0.

### Inverse sparse regression

If we knew *R* exactly, we could simply invert it and plug the inverse into (9). However, we only have access to the noisy estimate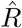, which is not necessarily well-conditioned or even invertible. Instead, we assume that the underlying directed graph is sparse. We observe that in (9), *G* is sparse if and only if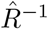 is sparse, and so we can view solving (9) as finding a sparse matrix inverse. We seek matrices *U, V* with *V U* = *I* that minimize the loss,

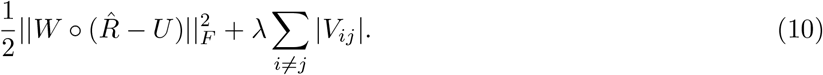

We minimize this loss using alternating direction method of multipliers (ADMM)^34^. Let Θ^*k*^ be a matrix of Lagrange multipliers. The updates for *U* ^*k*^, *V* ^*k*^ and Θ^*k*^ are

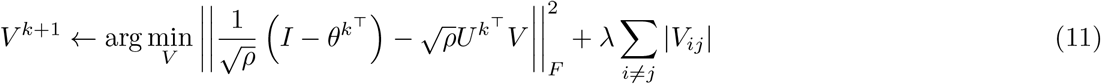

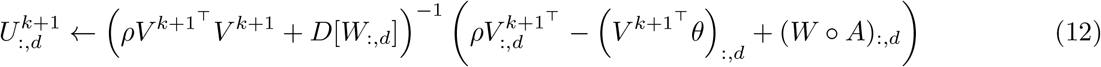

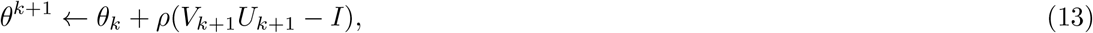

where *ρ* is the penalty parameter^34^. The update for *V* ^*k*+1^ is a straightforward LASSO regression while the update for *U* is a system of linear equations. In both cases we use an iterative solver and fit 10 iterations rather than running until convergence. We always start from the initial condition *U*_0_ = *V*_0_ = *I*. For the derivation of these equations including the specifics of how we tune the penalty parameter see the Supplemental note.

### The approximate DAG constraint

The general *inspre* model does not assume that the underlying graph *G* is a DAG. However, in some cases it may be preferable to fit a model that assumes a DAG, or you may obtain better convergence in practice by assuming a DAG. It is known that that *G* is a DAG if and only if *D*[(*I − G*)^*−*1^] = *I*^11^. Using 9, we have that

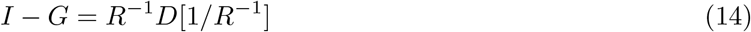

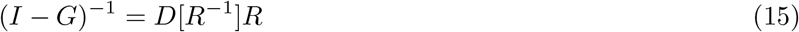

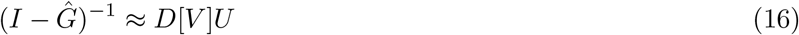

Thus, we can implement an approximate DAG constraint by constraining *D*[*V*] = *D*[*U*] = *I* in the above regression (10).

### Cross-validation and setting the LASSO penalty

We provide several cross-validation metrics to aid in selection of the LASSO penalty. We use 5-fold cross validation to calculate the loss

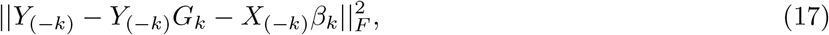

the loss

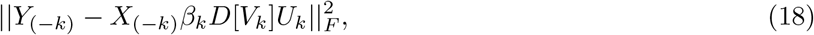

and the loss

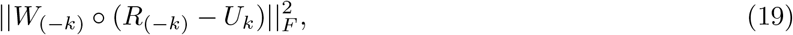

where *Y*_(*−k*)_, *X*_(*−k*)_ are the test observations for cross-validation iteration *k* and *G*_*k*_, *β*_*k*_ are the network and intervention effects estimated using the training observations *Y*_(*k*)_,*X*_(*k*)_. In the latter, we use the formulation from 2 with the approximation in 16.

In addition, we calculate stability estimates using the Stability Approach to Regularization Selection (StARS,^35^). StARS leverages the intuition that larger values of *λ* yield graphs that are more stable under random re-samplings of the input data to construct an interpretable quantity representing the average probability that each edge is included in the graph for each value of *λ*. Let *ϕ*_*λ*_ be a *D× D* matrix where entry *i, j* is the probability that each edge *i, j* is included in the graph for regularization parameter *λ*. We estimate *ϕ*_*λ*_ by using the graph estimates from each cross-validation iteration,

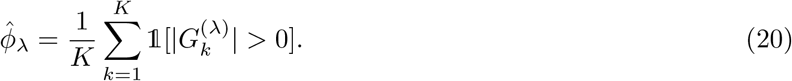

The instability measure *D*_*λ*_ is estimated as^35^

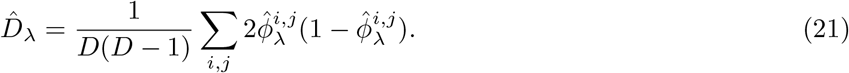

Clearly, *D*_*λ*_ = 0 for very large values of *λ*, where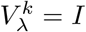 for every mask *k*. As *λ* becomes smaller, *D*_*λ*_ rises, but as *λ* approaches 0, *D →* 0 as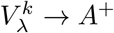. Following^35^, we first normalize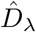 by setting it to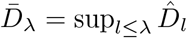 and then choose the smallest value of *λ* with stability below a cut point *b*,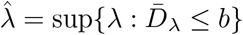.

In our simulation studies, we use the first loss and select the *λ* that minimizes the cross-validation error. In our analysis of the GWPS-data, this leads to underfitting due to high variability in the standard errors of the estimates of individual effects. Thus we choose *λ* based on analyzing each of these cross-validation metrics, aiming for a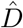 of around 0.01.

### Simulation details

We conducted extensive simulations according to the model 1 with *D* = 50 nodes while varying the density of the network, the network edge weights, and intervention effect sizes. We simulated random and scale-free networks, with and without cycles, and with and without confounding. We simulated 10 independent graphs for each of these 64 settings. To avoid issues of varsortability^36^, aid interpretability in setting simulation parameters, and more accurately represent common practices in genetics, all phenotypes are simulated with mean 0 and variance 1.

To vary the network density we varied the probability of edge inclusion (in random graphs) and the probability of adding an edge between existing nodes (in scale-free graphs) such that the average node degree in each graph was 2 (low-density) or 4 (high-density). To generate edge weights, we used a PERT distribution^37^. We used the parameter *v* to represent the median per-variance graph effect while setting the minimum to *v/*2 and the maximum to 2*v*. We used the setting *v* = 0.15 to represent weak effects and the setting *v* = 0.3 to represent strong effects. To set the intervention effect, we used a normalized effect setting equal to the number of standard deviations the mean of *Y*_*i*_ shifted given the presence of the intervention *X*_*i*_ = 1. For weak effects we used *β* =*−* 1 and for strong effects we used *β* =*−* 2. To generate confounding, we leveraged the observation that confounding effects are dense components in the graph structure^38^. We then simulated a specified number of additional unobserved full out-degree nodes. For edge weights, we again used a PERT distribution, but with the median effect set to *v*^*α*^, where *α* is the mean path length for that graph type: *α* = log(*D*) for random graphs^39^ and *α* = log(*D*)*/* log(log(*D*)) for scale-free graphs^40^. In each setting, we set *C* such that the mean variance in each phenotype explained by confounding factors was 10%.

We compared *inspre* against observational methods LiNGAM^14^, notears^11^, and golem^15^; and interventional methods GIES^16^, dotears^10^ and igsp^12^. For GIES and LINGAM, we used the implementation in the pcalg R package. We used default parameter settings for all methods with two exceptions: 1) for all methods, we used edge thresholding set to *v/*4, 2) for methods that used *L*_1_ regularization, we used 5-fold cross-validation with a logarithmically decreasing sequence of 10 *λ* values from 1 to 10^*−*6^ and report results from the value of *λ* that minimized the cross-validation error for each method.

### Genome-wide Perturb-seq analysis

We obtained normalized essential-scale Perturb-seq data for K562 cell lines generated in Replogle *et al*^9^ from https://plus.figshare.com/ndownloader/files/35773075. We used two stage least squares to estimate perturbation effects given the targeted gene intervention. We retained genes where the estimated per-variance effect-size of the intervention on the target gene was at least *−*0.75 standard deviations, and where there were at least 50 cells receiving a guide targeting that gene.

We used 5-fold cross-validation to select from an exponentially decreasing sequence of 10 *λ* values with *λ*_*max*_ set to the absolute value of the maximum off-diagonal element of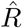 and *λ*_*min*_ to 0.1*×* this value (see above). We thresholded all elements of *Ĝ* with absolute value *<* 0.015 to 0, corresponding to roughly half the minimum FDR-5% significant ACE. We selected the *λ* value that minimized the cross-validation error. Due to improved convergence behavior, we use the approximate DAG constraint given above.

We calculated eigencentrality using the igraph R package function eigen centrality with default arguments. Note that this does not consider direction of edges in directed graphs. We conducted a beta regression of eigencentrality on 16 gene annotations using the betareg R package function betareg. We corrected for multiple testing using Holm’s method as implemented in the stats R package function p.adjust. For detailed descriptions and sources of each annotation see Table S7.

We obtained a list of genes that have been associated with 4 blood trait categories (platelet, reticulocyte, red blood cell, white blood cell) in genome-wide association studies from Morris et al^31^. We used the stats R package function t.test to test for a difference in mean path length between genes within each trait category versus mean path length between genes across trait categories. We corrected for multiple testing using Holm’s method as implemented in the stats R package function p.adjust. We used the stats R package function t.test to test for a difference in mean average effect of each gene on genes in each trait category against all genes not associated with any blood trait.

## Data and Code Availability

All code used in the production of this manuscript is available at https://github.com/brielin/inspre. GWPS data were obtained from FigShare https://plus.figshare.com/ndownloader/files/35773075. Full network results of our analysis are available on Zenodo at DOI 10.5281/zenodo.10002094.

## Acknowledgements

The authors would like to thank Júlia Domingo-Espinós and Mariia Minaeva for sharing curated geneannotation data. Funding for BCB is provided by NHGRI K99HG012373 and the Columbia Data Science Institute. Funding for DAK and BCB is provided by NIA U01AG068880. Funding for TL is provided by NIH R01AG057422 and NIMH R01MH106842. Funding for JM is provided by NHGRI K99HG012792.

## Competing Interests

Tuuli Lappalainen is a paid adviser or consultant of Variant Bio and GSK.

**Figure S1:**
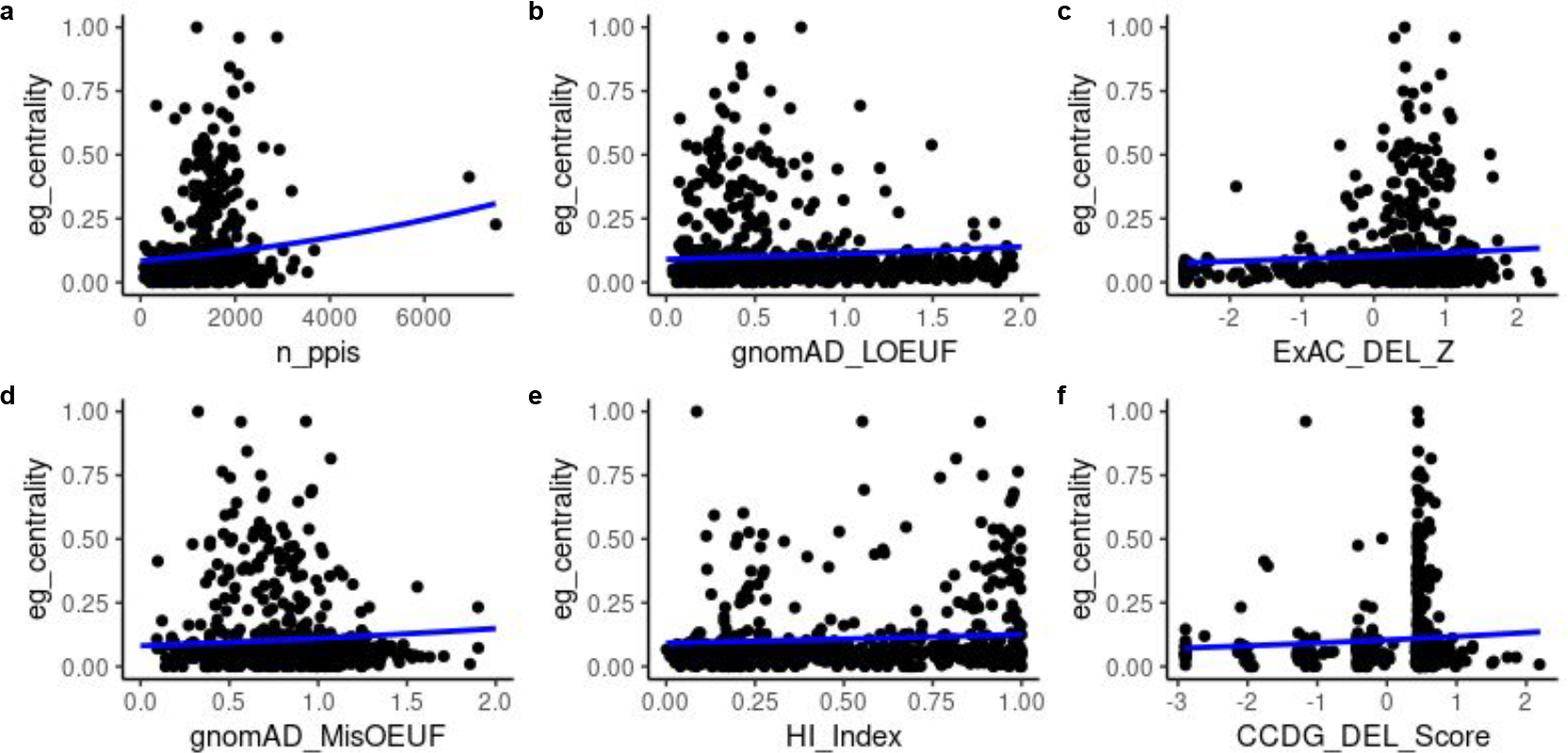
Eigencentrality vs gene-level annotation for associations in our analysis. Line represents beta regression fit.

**Figure S2:**
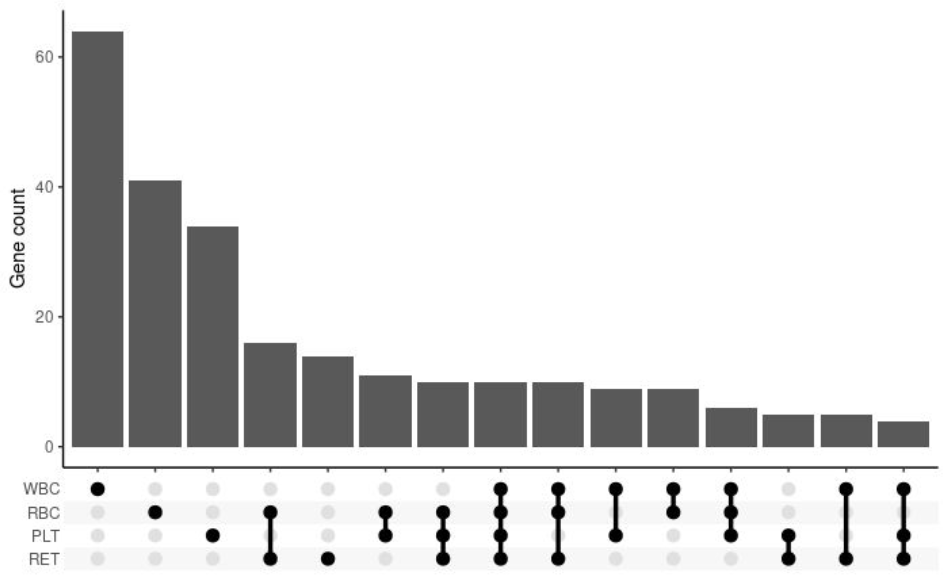
Upset plot of gene sets in our 788 analysis genes associated with blood traits in GWAS.

## Supplemental Note

### Alternating direction method of multipliers

First, consider the unweighted optimization problem

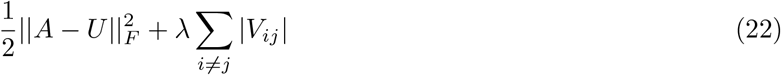

The augmented Lagrangian is,

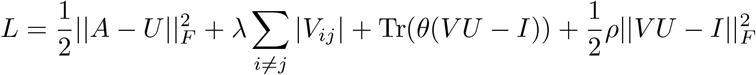

The update for *V* can be found by noticing that minimizing *L* is equivalent to solving a lasso regression with design matrix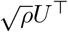 and response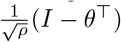,

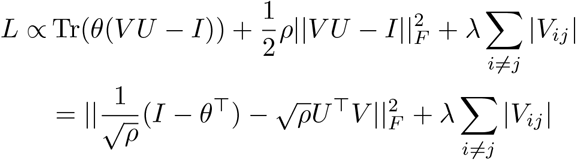

The update for *U* can be found by taking the gradient ▽_*U*_ *L* and setting it to 0,

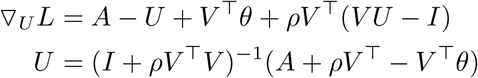

ADMM gives the update for *θ*^34^,

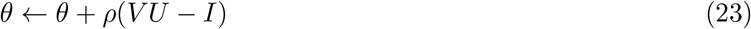

Now we consider the weighted version. Assume that in addition to the matrix *A*, we also have a matrix of standard errors of the entries of *A, S*_*A*_. Let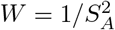 be a matrix of inverse variance weights. We now seek matrices *U, V* with *V U* = *I* that minimize the loss,

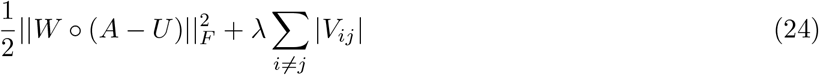

This does not effect the update for *V*, however the gradient of the augmented Lagrangian with respect to *U* is now,

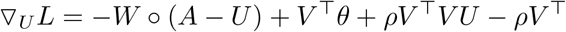

which separates over columns of *U*, giving the update

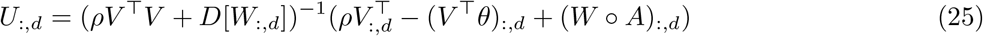

where here the *D* operator creates a matrix with *W*_:,*d*_ on the diagonal and 0 elsewhere.

ADMM also requires that we set the parameter *ρ*, which controls the balance in the objective between the primal and dual constraints^34^. We follow standard practice of setting rho to an initial value and increasing or decreasing it according to the ratio of the solution to the primal and dual feasibility constraints. The primal residual at iteration *k* + 1 is given by *r*^*k*+1^ = *V* ^*k*+1^*U* ^*k*+1^ *− I*. The dual residual is found by setting ▽_*U*_ *L*^*k*^ = 0 and evaluating it at *U*_*k*+1_

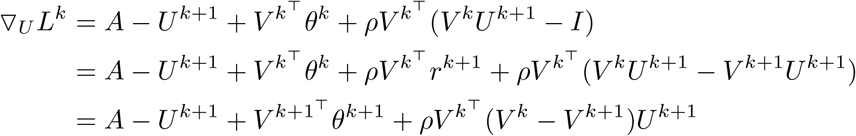

Therefore the dual residual is^34^

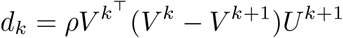

and we can adjust *ρ* as follows,

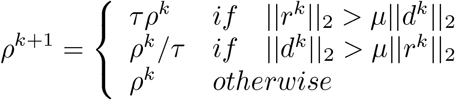

which reduces the impact of the initial choice of *ρ*. While this may appear to be a lot of parameters, they effect the convergence of the algorithm substantially more than the solution obtained. We always use the default values *ρ* = 10, *µ* = 10, *τ* = 2.

## References

1. Boyle, E. A., Li, Y. I. & Pritchard, J. K. An Expanded View of Complex Traits: From Polygenic to Omnigenic. Cell 169, 1177–1186 (2017).

2. Liu, X., Li, Y. I. & Pritchard, J. K. Trans Effects on Gene Expression Can Drive Omnigenic Inheritance. Cell 177, 1022–1034.e6 (2019).

3. Wray, N. R., Wijmenga, C., Sullivan, P. F., Yang, J. & Visscher, P. M. Common Disease Is More Complex Than Implied by the Core Gene Omnigenic Model. Cell 173, 1573–1580 (2018).

4. Parsana, P. et al. Addressing confounding artifacts in reconstruction of gene co-expression networks. Genome Biology 20, 4–9 (2019).

5. Verma, T. & Pearl, J. Equivalence and Synthesis of Causal Models in Proceedings of the Sixth Conference Annual Conference on Uncertainty in Artificial Intelligence (UAI-90) (1990), 220–227.

6. Chickering, D. M. Learning Bayesian Networks is NP-Complete, 121–130 (1996).

7. Hauser, A. & Ch, B. M. E. Characterization and Greedy Learning of Interventional Markov Equivalence Classes of Directed Acyclic Graphs Peter Bühlmann. Journal of Machine Learning Research 13, 2409–2464 (2012).

8. Angrist, J. D. & Imbens, G. W. Identification and Estimation of Local Average Treatment Effects (Feb. 1995).

9. Replogle, J. M. et al. Mapping information-rich genotype-phenotype landscapes with genome-scale Perturb-seq. Cell 185, 2559–2575.e28 (July 2022).

10. Xue, A., Rao, J., Sankararaman, S. & Pimentel, H. dotears: Scalable, consistent DAG estimation using observational and interventional data (May 2023).

11. Zheng, X., Aragam, B., Ravikumar, P. & Xing, E. P. DAGs with NO TEARS: Continuous Optimization for Structure Learning (2018).

12. Yang, K. D., Katcoff, A. & Uhler, C. Characterizing and Learning Equivalence Classes of Causal DAGs under Interventions. 35th International Conference on Machine Learning, ICML 2018 12, 8823–8839 (Feb. 2018).

13. Pachter, L. S. The network nonsense of Albert-LászlóBarabási — Bits of DNA

14. ShimizuShohei, O. H., HyvärinenAapo & KerminenAntti. A Linear Non-Gaussian Acyclic Model for Causal Discovery. The Journal of Machine Learning Research 7, 2003–2030 (Dec. 2006).

15. Ng, I., Ghassami, A. E. & Zhang, K. On the Role of Sparsity and DAG Constraints for Learning Linear DAGs. Advances in Neural Information Processing Systems 2020-Decem (June 2020).

16. Hauser, A. & Bühlmann, P. Characterization and Greedy Learning of Interventional Markov Equivalence Classes of Directed Acyclic Graphs. Journal of Machine Learning Research 13, 2409–2464 (Apr. 2011).

17. Erdős, P. & Rényi, A. On the evolution of random graphs. Publication ofthe Mathematical Institute of the Hungarian Academy ofSciences 5, 17–60 (1960).

18. Bollobas, B., Borgs, C., Chayes, J. & Riordan, O. Directed Scale-Free Graphs in Proceedings of the 14th Annual ACM-SIAM Symposium on Discrete Algorithms (SODA) (Jan. 2003), 132–139.

19. Bonacich, P. Power and Centrality: A Family of Measures. American Journal of Sociology 92, 1170–1182 (Mar. 1987).

20. Plaschka, C. et al. Architecture of the RNA polymerase II–Mediator core initiation complex. Nature 2015 518:7539 518, 376–380 (Feb. 2015).

21. Gamble, S. C. et al. Prohibitin, a protein downregulated by androgens, represses androgen receptor activity. Oncogene 2007 26:12 26, 1757–1768 (Sept. 2006).

22. Aitken, C. E. & Lorsch, J. R. A mechanistic overview of translation initiation in eukaryotes. Nature Structural & Molecular Biology 2012 19:6 19, 568–576 (June 2012).

23. Yotov, W. V. & St-Arnaud, R. Mapping of the human gene for the alpha-NAC/1.9.2 (NACA/1.9.2) transcriptional coactivator to Chromosome 12q23-24.1. Mammalian Genome 7, 163–164 (1996).

24. Karczewski, K. J. et al. The mutational constraint spectrum quantified from variation in 141,456 humans. Nature 581, 434–443 (May 2020).

25. Cassa, C. A. et al. Estimating the selective effects of heterozygous protein-truncating variants from human exome data. Nature genetics 49, 806–810 (May 2017).

26. Holm, S. A simple sequentially rejective multiple test procedure. Scandinavian journal of statistics, 65–70 (1979).

27. Wang, X. & Goldstein, D. B. ARTICLE Enhancer Domains Predict Gene Pathogenicity and Inform Gene Discovery in Complex Disease. The American Journal of Human Genetics 106, 215–233 (2020).

28. Bairoch, A. & Apweiler, R. The SWISS-PROT protein sequence database and its supplement TrEMBL in 2000. Nucleic Acids Research 28, 45 (Jan. 2000).

29. Huang, N., Lee, I., Marcotte, E. M. & Hurles, M. E. Characterising and predicting haploinsufficiency in the human genome. PLoS genetics 6, 1–11 (Oct. 2010).

30. Abel, H. J. et al. Mapping and characterization of structural variation in 17,795 human genomes. Nature 2020 583:7814 583, 83–89 (May 2020).

31. Morris, J. A. et al. Discovery of target genes and pathways at GWAS loci by pooled single-cell CRISPR screens. Science (New York, N.Y.) 380 (May 2023).

32. Sudlow, C. et al. UK Biobank: An Open Access Resource for Identifying the Causes of a Wide Range of Complex Diseases of Middle and Old Age. PLoS Medicine 12, 1–10 (2015).

33. Gasperini, M. et al. A Genome-wide Framework for Mapping Gene Regulation via Cellular Genetic Screens. Cell 176, 377–390.e19 (Jan. 2019).

34. Boyd, S., Parikh, N., Chu, E., Peleato, B. & Eckstein, J. Distributed optimization and statistical learning via the alternating direction method of multipliers. Foundations and Trends in Machine Learning 3, 1–122 (2010).

35. Liu, H., Roeder, K. & Wasserman, L. Stability approach to regularization selection (StARS) for high dimensional graphical models. Advances in Neural Information Processing Systems 23: 24th Annual Conference on Neural Information Processing Systems 2010, NIPS 2010, 1–14 (2010).

36. Reisach, A. G., Seiler, C. & Weichwald, S. Beware of the Simulated DAG! Causal Discovery Benchmarks May Be Easy To Game. Advances in Neural Information Processing Systems 33, 27772–27784 (Feb. 2021).

37. Clark, C. E. Letter to the Editor—The PERT Model for the Distribution of an Activity Time. 10.1287/opre.10.3.405 10, 405–406 (June 1962).

38. Chandrasekaran, V., Parrilo, P. A. & Willsky, A. S. LATENT VARIABLE GRAPHICAL MODEL SELECTION VIA CONVEX OPTIMIZATION 1. The Annals of Statistics 40, 1935–1967 (2012).

39. Fronczak, A., Fronczak, P. & Holyst, J. A. Average path length in random networks. Physical Review E - Statistical, Nonlinear, and Soft Matter Physics 70 (Dec. 2002).

40. Chen, F., Chen, Z., Wang, X. & Yuan, Z. The average path length of scale free networks (2006).

